# Actin-based myosin XXI (13) molecular motor is involved in early phase of *Leishmania* cytokinesis

**DOI:** 10.1101/2021.04.29.441917

**Authors:** Rani Bajaj, Chhitar M. Gupta

## Abstract

*Leishmania* genome encodes for two isoforms of myosin, but only Myosin XXI (Myo21), which is a novel form of myosin in that it contains two ubiquitin associated-like (UBA) domains towards the end of its tail structure, is expressed in both the promastigote and amastigote forms of this protozoan. Earlier studies have shown that in *Leishmania* promastigotes Myo21 besides localizing throughout the cell body and flagellum, it is prominently localized to the base of the flagellum. It has further been shown that this protein in the promastigotes plays an important role in regulating the cell morphology, motility, flagellum dynamics, growth and intracellular trafficking, As Myo21 depletion has been shown to result in reduced cell growth in culture, we considered it of interest to investigate whether the observed effect of Myo21 on the cell growth is mediated through its possible role in *Leishmania* cell division cycle. For this, we prepared heterozygous Myo21 mutants of *Leishmania* promastigotes (Myo21^+/−^cells) and then analyzed their morphology, growth and cell division cycle, using wild type *Leishmania* promastigotes (Myo21^+/+^ cells) as control. The cell division cycle was analyzed by employing flow cytometry and immunofluorescence microscopy. Flow cytometric analysis revealed that the G2/M to G1 phase transition in Myo21^+/−^ cell is significantly delayed, as compared to Myo21^+/+^ cells. Immunofluorescence confocal microscopic analysis indicated that Myo21^+/−^ cells encountered a significant delay in initiation of cytokinesis, which was mainly due to delay in the flagellar pocket division. Further analysis revealed that actin-based Myo21 motor is essentially required in the initiation phase of *Leishmania* cytokinesis.

## Introduction

The structural and regulatory elements that are essential for cell division have mostly been identified in higher eukaryotes from the Opisthokonta group, such as mammals, yeasts and nematodes. These basic and conserved elements broadly include cyclins and cyclin-dependent kinases that regulate the entry and exit of cell in distinct phases of the cell cycle, cellular motors that assemble microtubules for chromosomal segregation, and an actomyosin ring that drives the cytokinesis [1–4]. However, genomic sequencing of diverse eukaryotes revealed that gene encoding for proteins considered to be essentially involved in cytokinesis might not be conserved outside the Opisthokonta group. For instance, expression of non -muscle myosin II, which is known to produce force for cytokinesis, is restricted to the Opisthokonta group and is not present in other eukaryotic subgroups [5,6]. Thus, the molecular mechanisms involved in cytokinesis are perhaps more diverse as opposed to what was initially thought.

Trypanosomatids, including *Trypanosoma* and *Leishmania*, are a group of divergent model organisms whose genomes have already been sequenced [7,8] and for which genetic manipulation techniques are well established. These unicellular eukaryotic parasites have been widely studied to explore the molecular mechanisms that are involved in core processes, such as the cell division cycle. Amongst trypanosomatids, the cell division process of *T. brucei* procyclic form has been well characterized, which serves as a reference for *Leishmania* and other trypanosomatids, where only limited information is available. Events during cell division in trypanosomatids involve duplication and segregation of single copy organelles, such as the Golgi, the flagellum, the kinetoplast and the nucleus, and then cytokinesis generates two daughter cells [9]. However, *Leishmania* cell division differs from that of *T. brucei* in numerous ways. In *Leishmania*, the G2 phase is much shorter than that of *T. brucei*, and the order of the nucleus and kinetoplast division varies according to the species [10–12]. Besides this, in *Leishmania* promastigotes, the flagellum continuously grows from one cell cycle to the other and is accompanied by significant changes in cell morphology. The cells elongate during the G1 phase and then shrink and broaden during the G2/M phase, prior to cytokinesis [13]. A cleavage furrow forms along the longitudinal axis of the cell, which progresses from the anterior to the posterior end. However, little is known about the molecular events underlying promastigote cytokinesis. Our earlier studies have shown that in *Leishmania* promastigotes, actin dynamics plays a crucial role in early events, such as basal body and kinetoplast separation, flagellar pocket division and cleavage furrow progression, during their cell division [14]. As besides actin, these cells also express a novel form of myosin, myosin XXI [15], we considered it of interest to investigate whether myosin XXI, together with actin, is involved in *Leishmania* cytokinesis.

Although the *Leishmania* genome encodes for two isoforms of myosin, only one isoform belonging to a novel class of myosins, myosin XXI [6], which has later been reclassified as myosin 13 [16], is expressed in the *Leishmania* cells [15]. Myosin XXI (Myo21) is distributed throughout the cell body including the flagellum with prominent localization to the flagellum base [15], which is exclusively determined by its tail region [15]. Further, it has been shown that this protein in *Leishmania* promastigotes regulates cell morphology, flagellum assembly, cell growth and intracellular trafficking [17]. Here we report that Myo21 besides the above functions is also involved in *Leishmania* cytokinesis.

## Materials and Methods

### Antibodies

Anti-Myo21 and anti-actin antibodies were generated as described previously [18]. α-tubulin mouse monoclonal antibody (B-5-1-2; Cat.No.23948) was purchased from Santa Cruz Biotechnology company, whereas β-tubulin (Cat. No.T7816) mouse monoclonal antibodies were procured from Invitrogen. Secondary Alexa Fluor antibodies and HRP conjugated secondary antibodies were procured from Life technologies.

### *Leishmania* culture and growth analysis

The laboratory strain of *Leishmania* donovani (Dd8) was routinely maintained in high glucose Dulbecco’s modified Eagle’s medium (DMEM; Gibco, Life Technologies) supplemented with 10% of heat-inactivated foetal bovine serum (FBS; MP Biomedicals) with 40 mg L^−1^ gentamycin at 25°C. *Leishmania* promastigotes were transfected by electroporation, as described earlier [19]. Myo21 heterozygous mutants were maintained in the same medium with 10 μg ml^−1^ hygromycin B (Invitrogen). Episomally complemented heterozygous Myo21 mutants were maintained in the same medium with 10 μg ml^−1^ of hygromycin B and 10 μg ml^−1^ of G418 Sulfate (Sigma). For growth analysis, cells were inoculated at 10^5^ cells ml^−1^ density in the culture medium without antibiotic, and the cell number was counted every 24 hours for up to 15 days with a haemocytometer in three independent experiments.

### Generation of Myo21 heterozygous mutants and episomal complementation

The upstream flanking region of the Myo21 gene (5’FLK) was PCR amplified with primers F1 & R1. Similarly, the downstream flanking region of the Myo21 gene (3’FLK) was PCR amplified using primers F2 & R2. Hygromycin resistant gene was PCR amplified using F3 & R3 primers. Each of these fragments was cloned in the TZ57R vector to generate a 5’ - 5’FLK-Hyg-3’FLK - 3’ cassette and its release were confirmed by XbaI-BamHI digestion. Sequence verified clone was transfected in the *Leishmania* promastigotes by electroporation. Mutants were grown in the presence of hygromycin (10 μg ml^−1^) on DMEM agar plates.

Full-length Myo21 was PCR amplified (primers F4 and R4) and cloned in frame with C-terminus GFP tag in the pXG-GFP+ vector at BamHI site. For episomal complementation Myo21-GFP vector was transfected in Myo21^+/−^ cells and selected in the presence of 10 μg ml^−1^ hygromycin B and 50 μg ml^−1^ of G418 sulfate. Please refer the Table S1 for primer sequences.

### RNA isolation and qPCR

The fold change in expression of Myo21 mRNA in Myo21^+/−^ cells was determined using real-time PCR (qPCR). For this, total RNA was isolated from Myo21^+/+^ and Myo21^+/−^ promastigotes by TRIzol reagent (Ambion, Life Technologies) and DNase treated RNA was reverse transcribed with MMLV reverse transcriptase (NEB). Primers F5-R5, F6-R6 and F7-R7 were used to amplify the regions of *Leishmania donovani* actin, tubulin and Myo21 gene, respectively. Each cDNA sample was amplified using SYBR Green (Kapa Biosystems) on a fast real-time PCR system (Applied Biosystems). Results obtained were quantified by the delta-delta CT method with actin as a reference gene and tubulin as an endogenous control to normalize each sample. All qPCR experiments were performed using SYBR green at 95°C for 20 sec, 55°C for 30 sec and 72°C for 1 sec for 40 cycles. The specificity of the reaction was verified by melt curve analysis. Please refer the Table S1 for primer sequences.

### Western blotting

*Leishmania* promastigotes from the mid-log phase were used for the expression analysis. Lysates were prepared by washing the cells with PBS (phosphate buffer saline, pH 7.4) and boiling the resuspended cells in an SDS-PAGE sample buffer. Protein estimation was done by Bradford assay. Equal protein concentrations or equivalent cells were loaded and resolved on 10% SDS–PAGE. Gels were electroblotted on nitrocellulose membrane and then the membrane was incubated in the blocking buffer (5% skimmed milk in TBS, pH 7.4). The membrane was treated with primary antibodies (anti-Myo21: 1:2500; anti-LdAct: 1:5000) diluted in blocking buffer. After washing off the unbound antibodies, the membrane was probed with HRP-conjugated secondary antibodies (HRP-conjugated anti-rabbit IgG or anti-chicken IgY; Invitrogen; 1:5000). To develop the blots chemiluminescence ECL substrate (Amersham, GE Healthcare) was used and images were captured in an imaging system from the Bio-Rad (Molecular Imager^®^ ChemiDoc™ XRS+). Band intensity was quantified by Gel Quant.net software.

### Immunofluorescence microscopy

Cells were washed two times with PBS and were adhered to poly–L–lysine coated coverslips for 10 min. Cells were fixed in 2% paraformaldehyde (w/v) for 30 minutes and were washed with 0.5% PBS–glycine (w/v). Fixed cells were treated with PBS containing 0.5% (v/v) Triton X-100 to permeabilize them for labeling with the antibodies, followed by blocking in PBS with 3% bovine serum albumin (BSA; w/v). The immunolabeling of cells was performed with primary antibodies: anti-Myo21 (1:500), anti-α/β tubulin (1:1000) and anti-LdAct (1:10,000); secondary antibodies: anti-rabbit Alexa Fluor 488, anti-mouse Alexa Fluor 568 and anti-chicken Alexa Fluor 568 antibodies. Coverslips were mounted using prolonged diamond anti-fade mounting media containing DAPI (4, 6-diamidino-2-phenylindole; Invitrogen) and imaging was done on Nikon laser scanning confocal microscope Plan Apo VC 100X (oil) lens.

Mid-log phase cells were stained with DAPI for nuclear and kinetoplast labeling. These cells were analyzed microscopically and categorized depending on the number of nuclei and kinetoplasts per cell into 1N1K, 2N2K, 2N1K, 2N2K and others. For quantification, experiments were performed in triplicates (n≥600) separately for wild type cells (Myo21^+/+^), heterozygous mutants (Myo21^+/−^) and episomally complemented mutant cells (Myo21^+/−comp^).

For flagellar pocket division analysis, cells were treated with ConA-rhodamine, as described earlier [14]. Cells were washed two times with ice-cold PBS and then fixed with paraformaldehyde (4%) at room temperature for 30 minutes. The fixed cells were rinsed thrice with the ice-cold PBS and then resuspended in 1 ml PBS containing ConA-rhodamine (1:500; 1mg ml^−1^ stock) and incubated at room temperature for 2 hours. Subsequently, excess of ConA-rhodamine was removed by washing the cells twice with the PBS before adhering them on poly-L-lysine coated glass coverslips. Coverslips were mounted using prolonged diamond anti-fade mounting media containing DAPI.

To adjust the gain/offset and blank the background signals for the image data collection, a negative control slide was used. Brightness/contrast was adjusted, when required images were cropped and panels of images were arranged in Adobe Photoshop for presentation.

### Cell cycle analysis

To perform cell cycle analysis, mid-log phase *Leishmania* promastigotes (≤10^7^ cells ml^−1^) were synchronized with HU (hydroxyurea) [14]. Cells were centrifuged at 1400 × g for 10 min at 4°C, followed by their suspension in the fresh DMEM with 200 μg ml^−1^ HU (Sigma) and incubated overnight (12-14 hours). After incubation, the cells were washed with the PBS twice and then suspended in the fresh DMEM media without HU. Cells were collected (~10^7^) every 2 hours for up to 12 hours. The cell aliquot was mixed with 150 μl of fixative solution (20mM sodium phosphate, 40mM citric acid, 200mM sucrose and 1% Triton X-100) and incubated for 5 minutes at room temperature. To the fixed cells, 350 μl of diluent buffer (125mM MgCl_2_ in PBS) was added and until further use, samples were stored at 4°C. To quantify the DNA, cells were incubated with 50 μg RNase (5 mg ml^−1^ in 0.2M sodium phosphate buffer, pH 7.0) for two hours at 37°C and then stained with 50 μg propidium iodide (5 mg ml^−1^ in 1.12% sodium citrate) for one hour at 25°C, followed by equilibration at 4°C for overnight. A Gallios flow cytometer (Beckman Coulter) was used to generate the data from the processed samples, and the ModFit software was used to calculate proportions of the G1, S and G2-M population. For each sample, 20,000 events were collected.

### Statistical Analysis

As and when required, results were expressed as the mean ±S.D. (standard deviation) from independent experiments performed in triplicates. Statistical analysis was done by ANOVA tests in Microsoft Excel. A p-value was considered significant if <0.05.

## Results

### Generation and validation of Myo21 heterozygous mutants

*Leishmania* is a diploid organism, where the Myo21 gene is present in a single copy with two alleles. It has earlier been reported that Myo21 null mutants do not survive in culture, which indicated that Myo21 expression is perhaps essential for the survival of the parasite [17]. Therefore, we generated heterozygous Myo21 mutants (Myo21^+/−^), where the Myo21 single gene allele was replaced with an antibiotic marker, i.e., hygromycin phosphotransferase (Hyg) gene, which confers resistance to hygromycin B.

Upstream and downstream flanking regions of the Myo21 gene was PCR amplified and cloned at 5’ and 3’ ends of hygromycin resistant gene to generate Hyg-cassette (see the materials and methods; Fig. S2). Sequence verified Hyg-cassette was transfected in *L. donovani* promastigotes by electroporation and hygromycin resistant transfectants were selected on DMEM-agar plates in the presence of hygromycin B (10 μg ml^−1^). The levels of Myo21 transcript and protein in heterozygous mutants were analyzed by qPCR and western blotting, respectively. The analysis revealed that as compared to the Myo21^+/+^ cells, transcripts and protein levels of Myo21 were reduced by about 70% in Myo21^+/−^ cells (Fig. S3). For add back analyses, Myo21^+/−^ mutants were complemented by episomal expression of the Myo21-GFP gene (Myo21^+/−comp^).

### Morphology and growth analysis of Myo21^+/−^ cells

Earlier studies from our group have shown that depletion of the Myo21 intracellular pool adversely affected the cell growth, morphology, motility and flagellar length [17]. The Myo21 depleted cells became stumpy and nonmotile and their flagellum length was significantly reduced [17]. To reconfirm these observations, we performed confocal microscopy analysis of the mutant parasites for intracellular distribution of Myo21 as well as for cell morphology. As expected, Myo21 heterozygous mutants appeared stumpy and short with substantially reduced flagellar lengths, where Myo21 showed the enriched presence in a dot-like pattern at the proximal region of the flagellum (Fig.1). The growth analysis revealed reduced growth of the mutants, as compared to wild type cells, which was restored close to normal in Myo21^+/−comp^ cells (Fig. 2A).

**Fig. 1.**
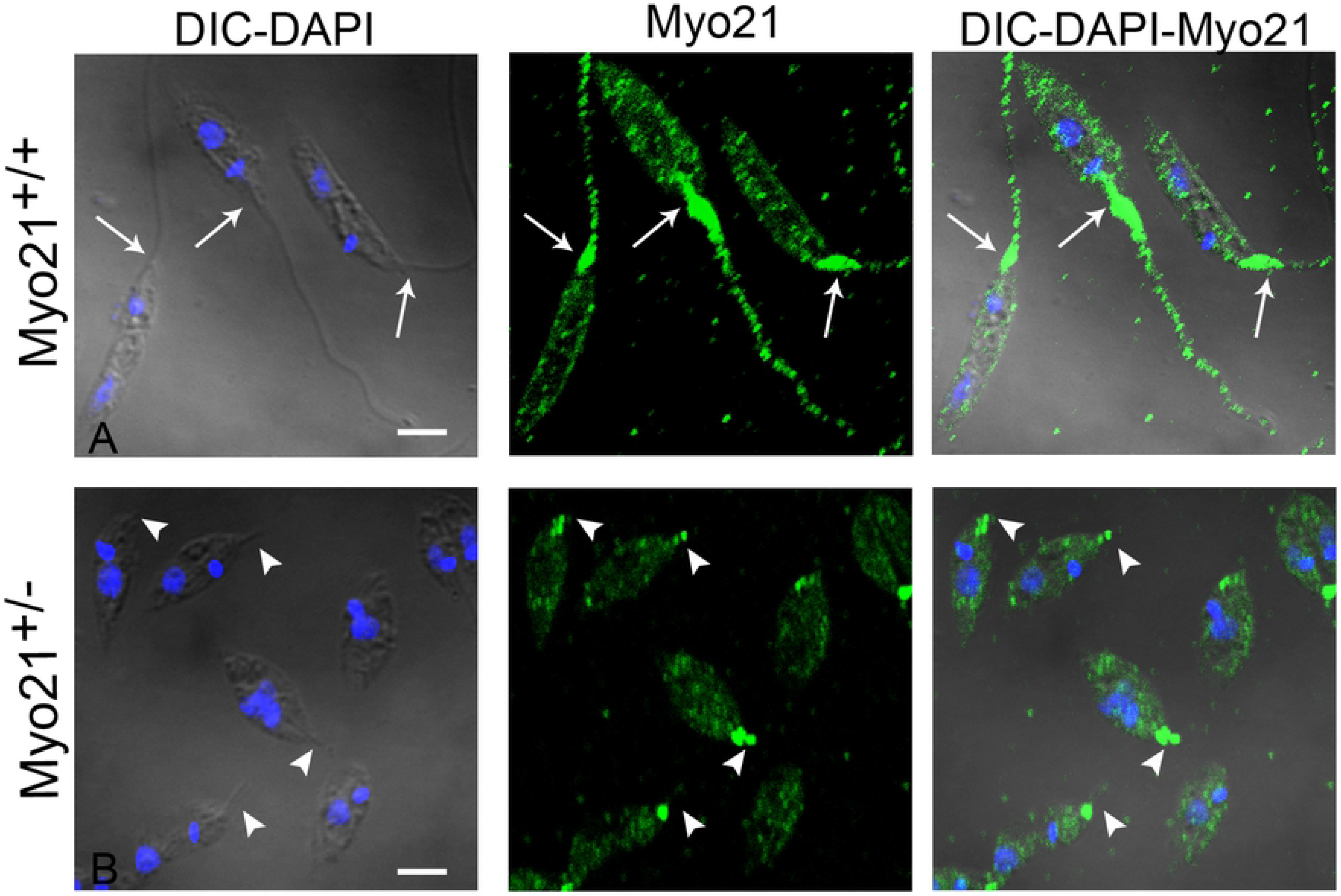
Intracellular distribution of Myo21 determined by immunofluorescence microscopy. Immunofluorescence images of (A) Myo21^+/+^ and (B) Myo21^+/−^ cells labeled with anti-Myo21 antibodies (green) showing localization of the Myo21. Scale bar-2 μm. Normal flagellar length in Myo21^+/+^ cells is marked with the arrows. Myo21^+/−^ cells are short and stumpy with reduced length of the flagellum, marked by the arrowheads.

**Fig. 2.**
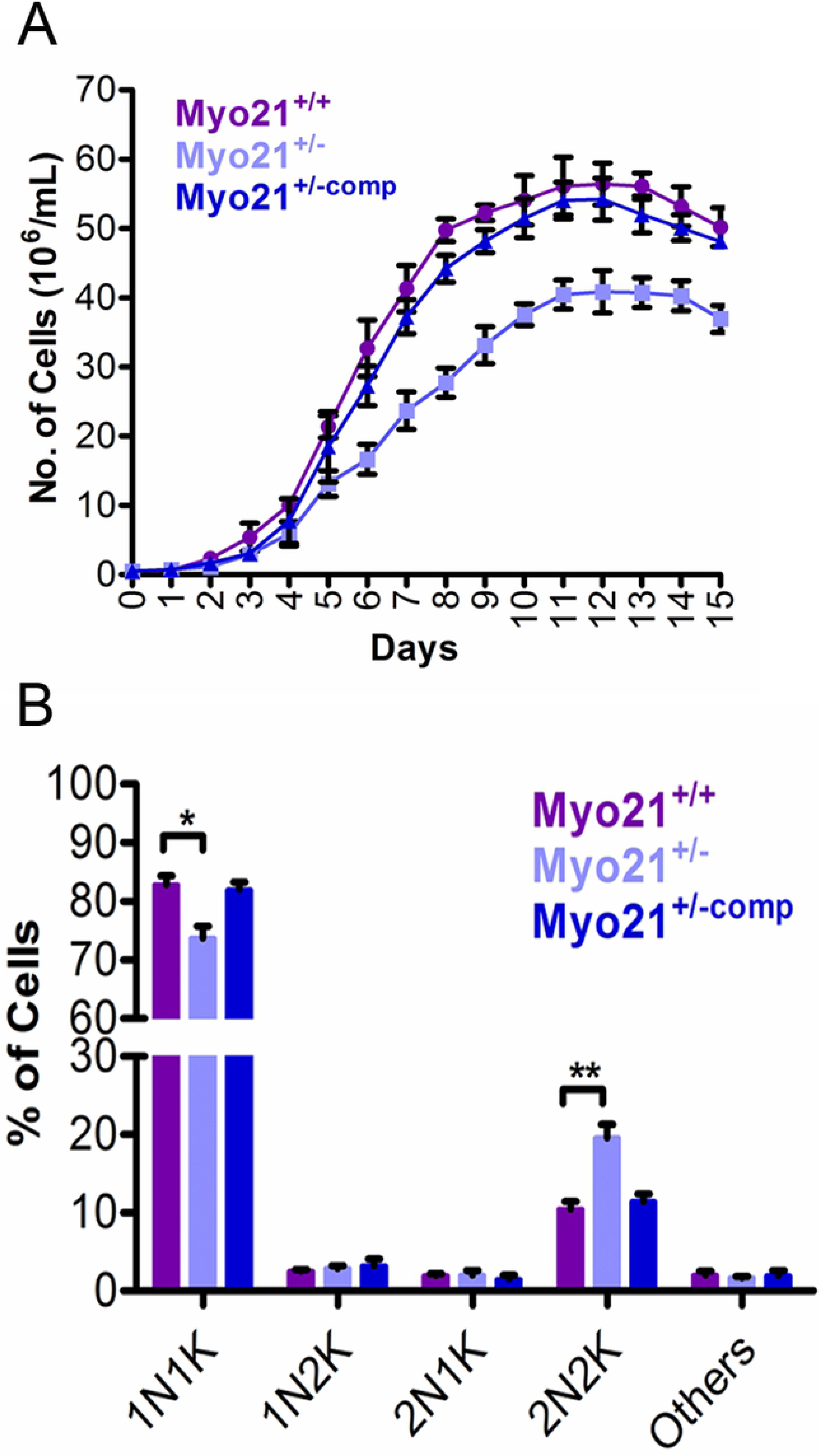
Growth analysis of Myo21 heterozygous mutants. (A) Myo21^+/+^ cells (purple circle), Myo21^+/−^ cells (light blue square) and Myo21^+/− comp^ cells (dark blue triangle) were seeded each at an initial cell density of about 10^5^ cells ml^−1^ and were grown in DMEM containing 10% FBS without antibiotics. The cell numbers were counted every 24 hours for up to 15 days with a hemocytometer. Myo21^+/−^ cells showed a significant reduction in their growth, compared to Myo21^+/+^ cells. However, the growth was restored to normal in Myo21^+/−comp^ cells. The data presented are the mean of three independent experiments ± S.D. (B) Cell division analysis in Myo21^+/−^ cells based on the nucleus and kinetoplast configuration. Mid-log phase Myo21^+/+^ (purple bars), Myo21^+/–^ (light blue bars) and Myo21^+/–comp^ (dark blue bars) cells were stained with DAPI to label their nucleus (N) and kinetoplast (K). Quantification of the nucleus and the kinetoplast numbers in each cell type was then carried out by fluorescence microscopic examination and accordingly cells were categorized as shown in the bar diagram. 1N1K, cells with one nucleus and one kinetoplast; 1N2K, cells with one nucleus and two kinetoplasts; 2N1K, cells with two nuclei and one kinetoplast and 2N2K, cells with two nuclei and two kinetoplasts. In each experiment, about 200 cells were analyzed. The mean values from three independent experiments were calculated and shown here with ±S.D. p-values being 0.02, 0.009 for 1N1K and 2N2K, respectively. Statistical analysis was done by ANOVA test and a p-value was considered significant if <0.05.

### Depleted level of Myo21 results in impaired cell division

A detailed microscopic examination of the mid-log phase Myo21^+/+^, Myo21^+/−^ and Myo21^+/−comp^ cells was carried out after staining them with DAPI to label their nucleus and kinetoplast DNA. About 600 cells each of Myo21^+/+^, Myo21^+/−^ and Myo21^+/−comp^ were analyzed in three independent experiments. It was found that cultures of Myo21^+/−^ cells contained a significantly higher number of dividing cells with 2 nuclei and 2 kinetoplasts, as compared to Myo21^+/+^ and Myo21^+/−comp^ cells (Fig. 2B). These results indicated that the observed slower growth could be due to impaired cell division in Myo21^+/−^ cells.

### A prolonged G2/M phase in Myo21^+/−^ cells causes the accumulation of 2N2K cells

The observed slower growth and accumulation of dividing cells in Myo21^+/−^ cells could be due to a modified cell cycle progression. To investigate this possibility, the mid-log phase Myo21^+/+^, Myo21^+/−^ and Myo21^+/−comp^ cells were synchronized at the G1/S phase by hydroxyurea (HU, 200 μg ml^−1^) treatment for 12 hours. After the HU block was released, aliquots of the cells were collected at 2 hours interval up to 12 hours and stained with PI (propidium iodide) for analysis of cell cycle phases by flow cytometry (see the materials and methods). About 86% Myo21^+/+^, Myo21^+/−^ and Myo21^+/−comp^ cells were in the G1-S phase after the HU treatment.

The peak for the G2/M phase reached the maximum at the sixth hour in both Myo21^+/+^ (56.1±2.3%; n=3) and Myo21^+/−^ cells (56.61±6%; n=3), showing a similar pace of progression from G1 to S and S to G2/M phase, in both the cell types. At the eighth hour, the number of Myo21^+/+^ cells decreased in the G2/M phase (37.1±7.5%; n=3) with an increase in the number of G1 phase cells (43.8±6%; n=3). However, Myo21^+/−^ cells took significantly more time to progress from G2/M to the G1 phase (Fig. 3 & S4). In these cells, the reappearance of the G1 phase (46.2±1%; n=3) occurred at the twelfth hour, indicating a substantial delay in the G2/M phase. The Myo21^+/−comp^ cells navigated through the cell cycle phases similar to that of Myo21^+/+^ cells. These results established the role of Myo21 in the G2/M phase of the cell division cycle in *Leishmania*.

**Fig. 3.**
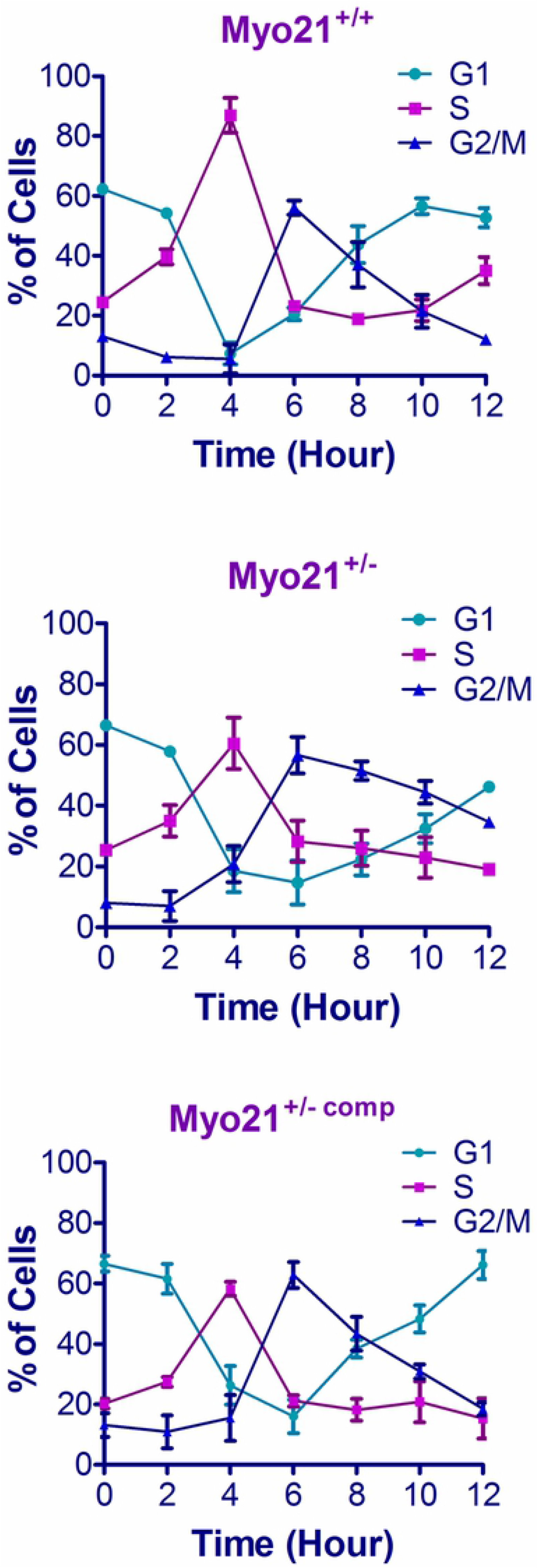
Graphical representation of cell cycle progression data obtained from flow cytometry for Myo21^+/+^, Myo21^+/−^ and Myo21^+/−comp^ *Leishmania* promastigotes. Each cell type was synchronized at the G1/S phase border by hydroxyurea (HU) treatment and to follow cell cycle progression, cell aliquots were collected at a regular time interval of 2 hours up to 12 hours. Processed samples were analyzed by flow cytometry. Percentages of cells in G1 (light blue), S (purple) and G2/M (dark blue) phases were plotted on the Y-axis against the time on the X-axis. The values plotted are from three independent experiments ± S.D. Myo21^+/+^ and Myo21^+/−comp^ cells showed G2/M phase maxima at the sixth hour and re-entered into the G1 phase at the eighth hour. Although in Myo21^+/−^ cells the G2/M phase started at the sixth hour, the G1 phase reappeared only at the twelfth hour. Higher percentages of Myo21^+/−^ cells in the G2/M phase for a prolonged time, indicated a delayed progression through the G2/M phase.

### Myo21 depletion delayed cytokinesis in *Leishmania* promastigotes

Flow cytometry data revealed that by the eighth hour Myo21^+/+^ cells reappeared in the G1 phase, but Myo21^+/−^ cells at this time were largely arrested at the G2/M phase. To determine whether the arrest at the G2/M phase was due to delayed mitosis or cytokinesis, cells were labeled with DAPI after 8 hours of HU block release and then investigated for the nucleus and kinetoplast configuration. Immunofluorescence microscopy analysis revealed an increased frequency of dividing cells in Myo21^+/−^ cells having 2N2K configuration (42%, n=90), as compared to Myo21^+/+^ cells (22%, n=90) and Myo21^+/−comp^ cells (24.5%, n=66). Microtubule septum was clearly visible between the dividing 2N2K Myo21^+/−^ cells, indicating an arrest of these cells at the cytokinesis stage (Fig. 4).

**Fig. 4.**
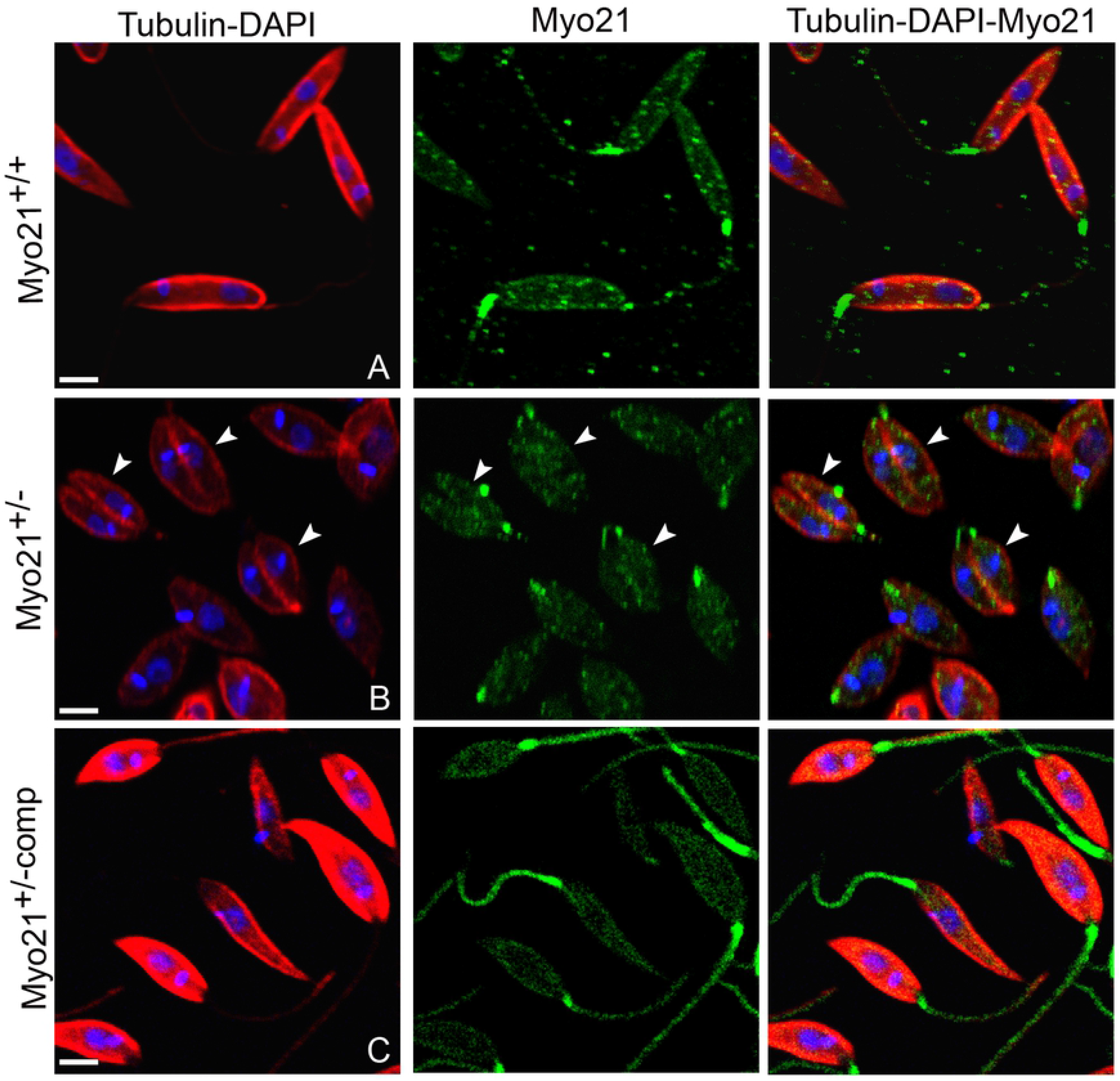
Immunofluorescence microscopic analysis of the synchronized Myo21^+/−^ cells in the division phase. Cells were synchronized by overnight treatment with hydroxyurea (HU), and after removing the HU block, cells were aliquoted, fixed and processed for imaging at the eighth hour. Cells were labeled for Myo21 (green), tubulin (red) and DNA (blue). Scale bar-2 μm. The number of cells analyzed for Myo21^+/+^: 90, Myo21^+/−^: 90 and Myo21^+/−comp^: 66 in three independent experiments. At the eighth hour, when most of Myo21^+/+^ and Myo21^+/−comp^ cells completed their division cycle, Myo21^+/−^ cells were arrested in the cytokinesis (cell separation) phase. Representative immunofluorescence images showing a higher number of dividing Myo21^+/−^ cells, compared to Myo21^+/+^ and Myo21^+/−comp^ cells. The arrowheads indicate Myo21^+/−^ cells with visible microtubule septa, indicating cells stuck in the final stage of the cell division (2N2K).

### Myo21, together with actin is involved in cytokinesis of *Leishmania*

Our prior studies have shown that actin dynamics are involved at least in the early phase of *Leishmania* cytokinesis [14], which prompted us to analyze the distribution of actin and Myo21 in the dividing *Leishmania* cells. Interestingly, about 80% (n=50) of dividing Myo21^+/+^ *Leishmania* cells showed enriched co-distribution of actin-Myo21 in the anterior region of the cell (Fig. 5). Further analysis of Myo21^+/+^ *Leishmania* cells revealed that enrichment of Myo21 along with actin at the anterior region occurs during the early stage of cytokinesis when division furrow ingresses (Fig. 6). Whereas, such co-distribution of the Myo21 and actin at the anterior region was not observed in the Myo21^+/−^ cells (n=18; Fig. 7). Together, these observations suggested that Myo21 and actin are involved in the early stages of *Leishmania* cytokinesis.

**Fig. 5.**
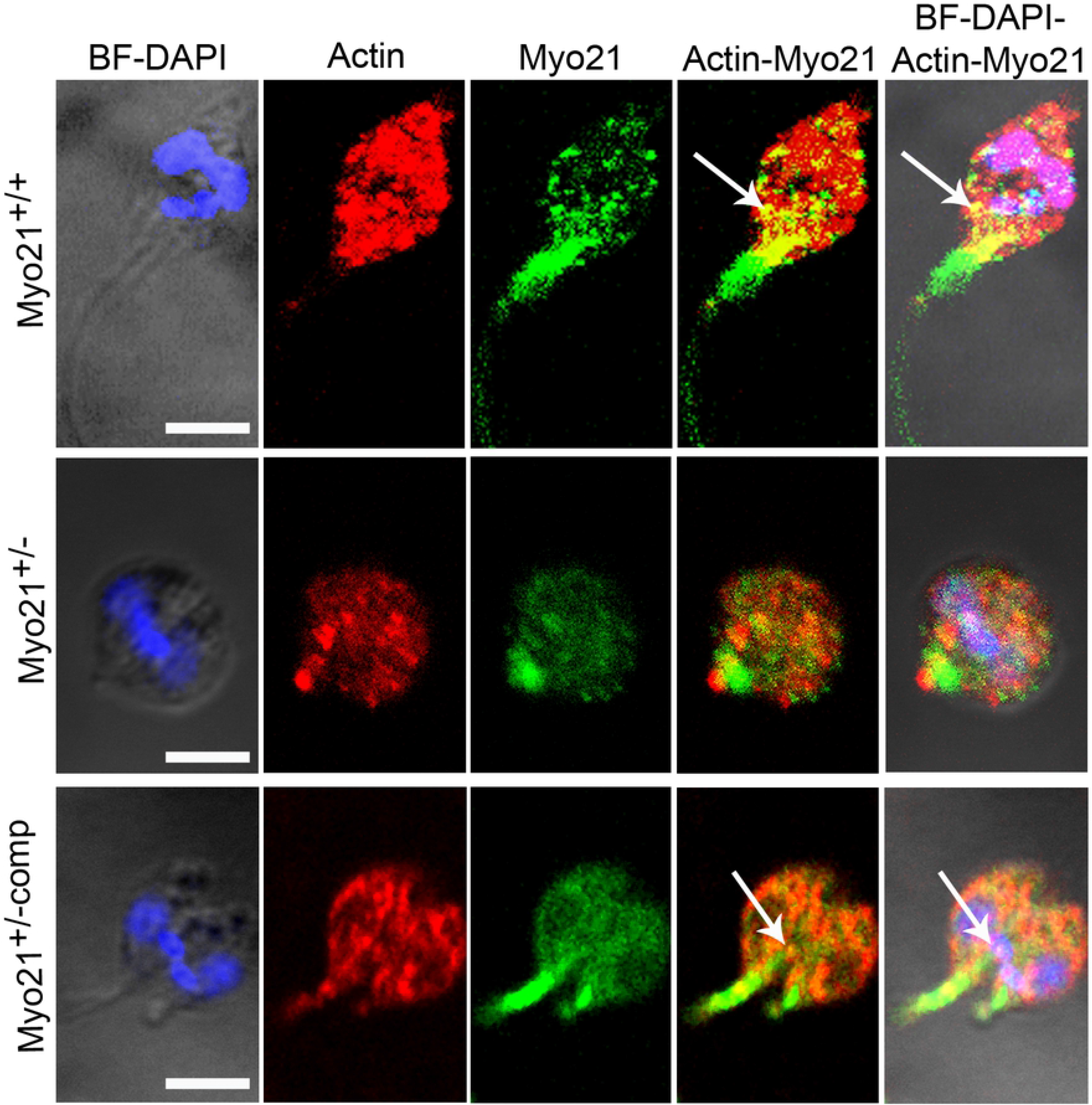
Immunofluorescence microscopic analysis of co-distribution of Myo21 and actin in dividing Myo21^+/+^, Myo21^+/−^ and Myo21^+/−comp^ *Leishmania* cells. Cells were labeled with the anti-Myo21 antibodies (green) and anti-actin antibodies (red) and mounted in DAPI to label the nuclear and kinetoplast DNA (blue). BF-Brightfield. Scale bar-2 μm. The number of cells analyzed for Myo21^+/+^: 50, Myo21^+/−^: 18 and Myo21^+/−comp^: 12 in three independent experiments. During the early stage of cytokinesis Myo21 co-localized with actin at the anterior end of the Myo21^+/+^ and Myo21^+/−comp^ *Leishmania* cells, as indicated by the arrow.

**Fig. 6.**
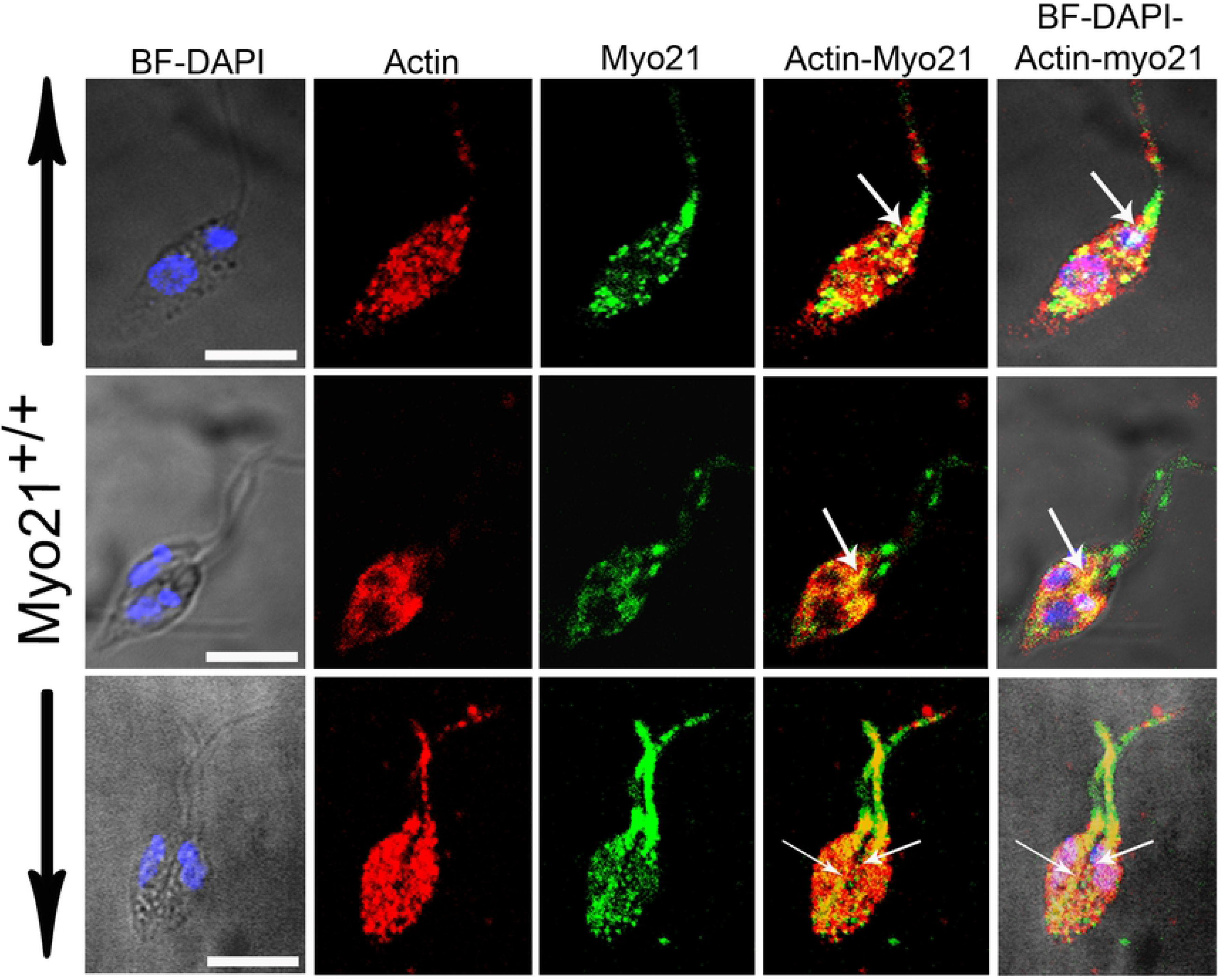
Intracellular distribution of the Myo21 and actin in Myo21^+/+^ *Leishmania* cells analyzed by confocal microscopy. Wild type *Leishmania* promastigotes were labeled with anti-Myo21 antibodies (green) and anti-actin antibodies (red) and mounted in DAPI to label the nuclear and kinetoplast DNA (blue). BF-Brightfield. Scale bar-5μm. About 50 cells were analyzed in three independent experiments. During the early stage of cytokinesis Myo21 co-localized with actin mainly at the anterior end of the cell. The arrow marks co-localization of actin and Myo21.

**Fig. 7.**
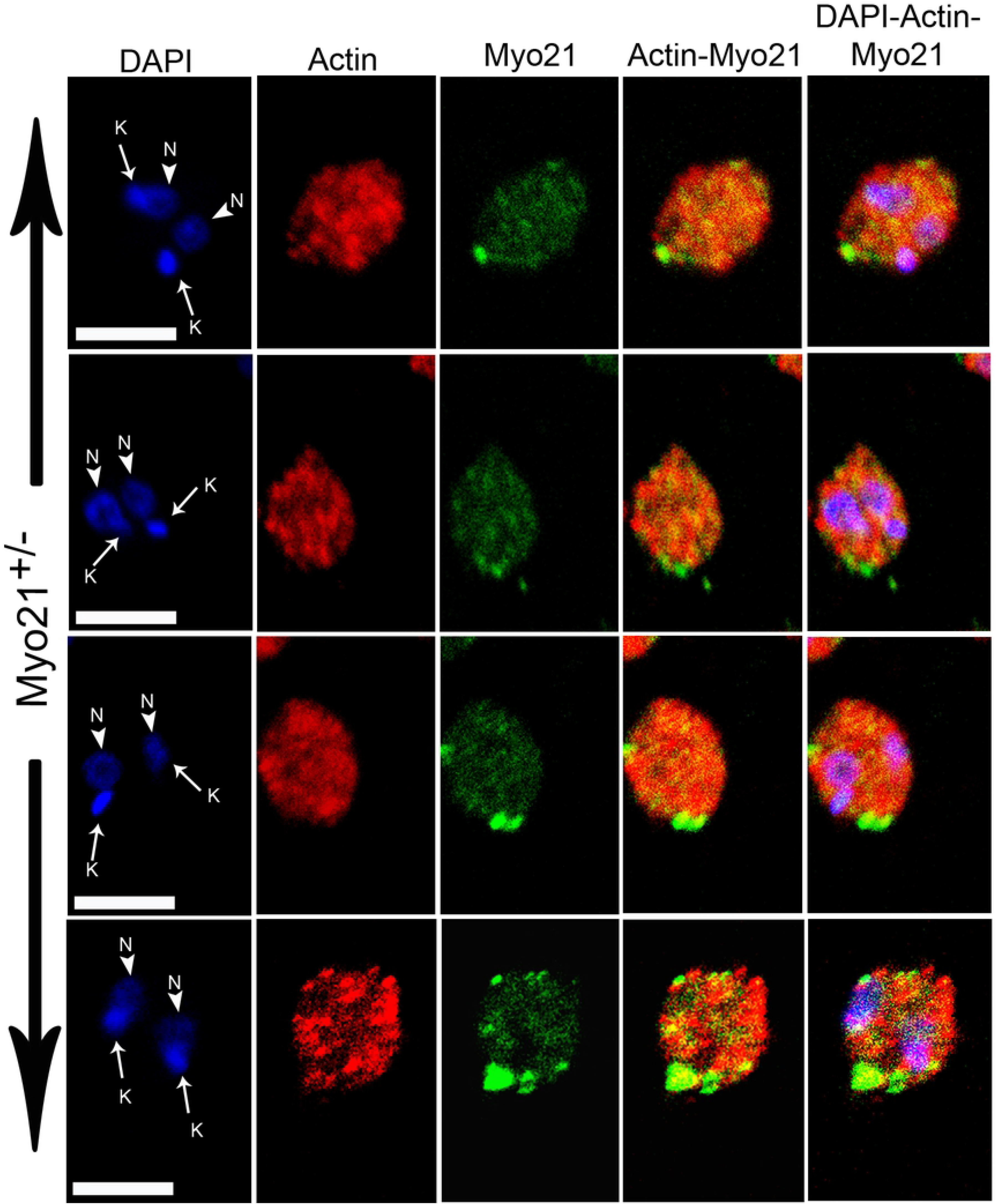
Intracellular distribution of the Myo21 and actin in Myo21^+/−^ *Leishmania* cells as analyzed by confocal microscopy. Heterozygous Myo21 mutants were labeled with anti-Myo21 antibodies (green) and anti-actin antibodies (red). Mounting was done in DAPI to label the nucleus (N) and kinetoplast (K). BF-Brightfield. Scale bar-2μm. 18 cells were analyzed in three independent experiments.

### Delayed furrow progression in Myo21^+/−^ cells is caused by delayed flagellar pocket division

The cell cycle progression in trypanosomatids is coupled with the assembly of the new flagellum and flagellar pocket division [20,21]. The events upstream of the furrow ingression during *Leishmania* cytokinesis should therefore include the assembly of the new flagellum and flagellar pocket division [9]. The flagellar pocket begins to divide before the onset of mitosis, wherein the pocket membrane invaginates adjacent to the kinetoplast and progresses towards the cell surface to split into two [9]. As our prior studies have shown that cell division cycle in ADF (actin polymerizing factor) knockouts is adversely affected due to delayed cytokinesis caused by delayed flagellar pocket division in *Leishmania* cells [14], we speculated that like in ADF depleted cells, delayed cytokinesis in Myo21 depleted cells might have been caused by delayed flagellar pocket division. To confirm this conclusion, we labeled the G2/M phase cells with ConA rhodamine, which is known to selectively label the flagellar pocket membrane [22], and then examined the cells under a fluorescent microscope (see the material and method). About 78% (77.5±4.1%; n=36) of Myo21^+/+^ cells and 74% (73.8±6.7%; n=12) of Myo21^+/−comp^ cells exhibited two well segregated flagellar pockets situated between the two dividing cells, whereas in Myo21^+/−^ cells, only about 47% (46.5±3.5 %; n=41) of dividing cells had separated flagellar pockets and the remaining cells (53.5±3.5 %; n=41) displayed a single flagellar pocket between the dividing cells (Fig. 8). These results clearly indicate that Myo21 together with actin regulates cytokinesis, especially flagellar pocket division, in *Leishmania* promastigotes.

**Fig. 8.**
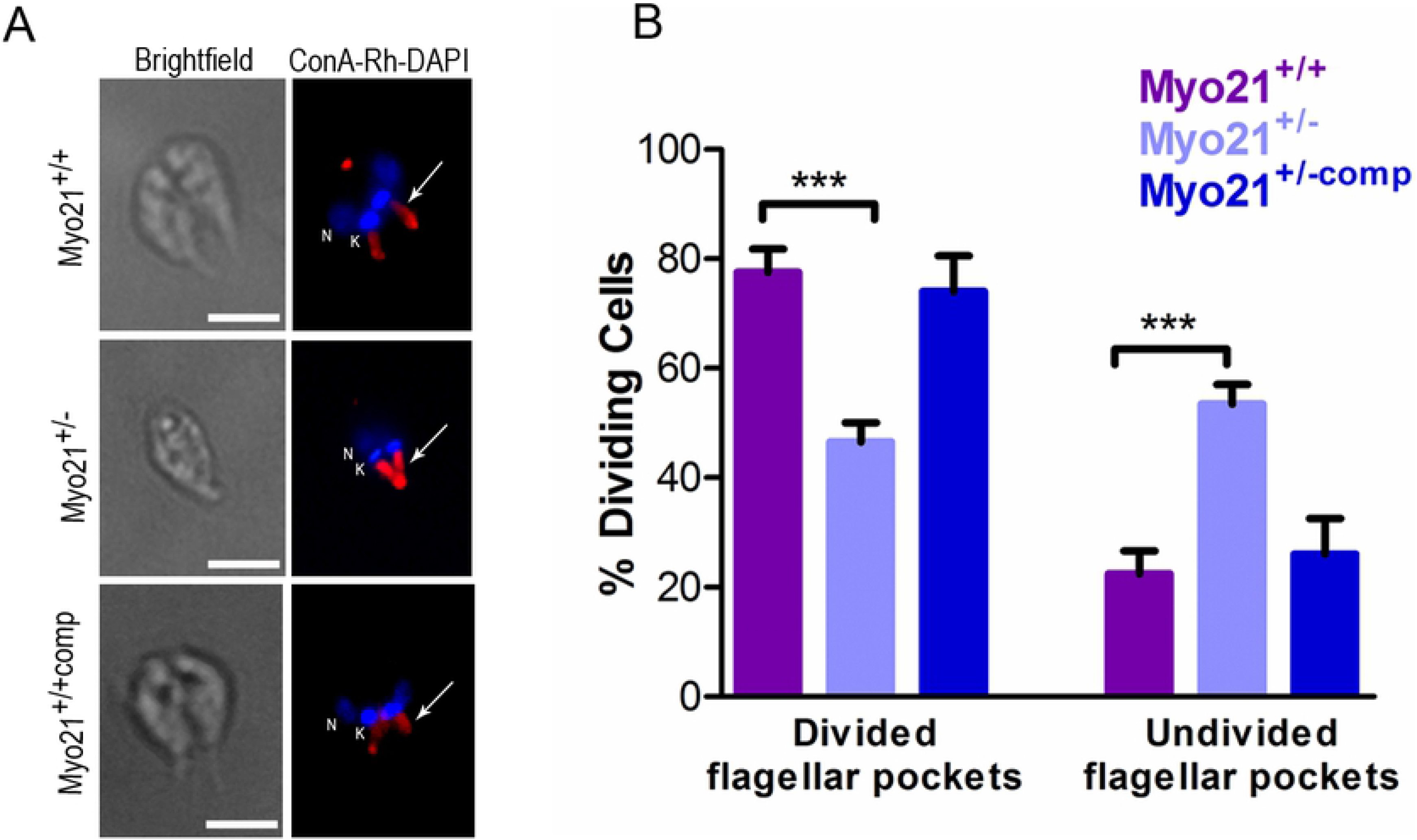
Analysis of flagellar pocket division. (A) Fixed cells were labeled with ConA-rhodamine (ConA-Rh; red) to label the flagellar pockets. Nuclei (N) and kinetoplasts (K) were stained with DAPI (blue). Epifluorescence microscopy images showing flagellar pockets divided into two in the Myo21^+/+^ and Myo21^+/−comp^ cells compared with an undivided flagellar pocket in Myo21^+/−^ cell. In dividing cells there are two nuclei and two kinetoplasts. The arrow marks the flagellar pockets. Scale bar-10 μm. (B) Quantitative analysis of flagellar pocket status. For each cell type dividing cells were counted after labeling the flagellar pockets with the ConA-rhodamine. Dividing cells with the two well-separated pockets were categorized as divided flagellar pockets and those with a single flagellar pocket were counted as undivided pockets. The number of cells analyzed for Myo21^+/+^: 36, Myo21^+/−^: 41 and Myo21^+/−comp^: 12 in three independent experiments. Statistical analysis was done by the ANOVA test and a p-value was considered significant if ≤0.05. P*** being 0.0006.

## Discussion

In the present study, we have demonstrated that the reduced growth of Myo21^+/−^ cells is due to the accumulation of the dividing cells, which in turn resulted from a delayed G2/M phase progression during the cell division cycle. Myo21^+/−^ *Leishmania* cell cycle proceeded normally till completion of karyokinesis but thereafter resulted in accumulation of the 2N2K cells, a signature of cytokinesis arrest. Despite the formation of the microtubule septum, Myo21^+/−^ 2N2K cells were unable to further divide into two daughter cells, suggesting the arrest at the furrow-ingression stage. Detailed microscopic analysis showed a cell cycle-dependent redistribution of Myo21 and actin in *Leishmania* cells. Further, it showed that the delayed furrow ingression in Myo21 depleted cells was linked to a delayed flagellar pocket division. It may therefore be suggested that Myo21, together with actin is required for separation of the daughter cells.

The flagellum emerges from the flagellar pocket; a specialized region formed by the invagination of the plasma membrane and is the sole site where endocytosis and exocytosis occur in the trypanosomatids [23,24]. The flagellar pocket is devoid of subpellicular microtubules and thus it is likely that the actin cytoskeleton manages dynamic activities of this region, which involve vesicular trafficking and furrow formation during cytokinesis. Indeed, actin dynamics has been implicated in the flagellar pocket division and the furrow ingression during cytokinesis in *Leishmania* promastigotes [14].

Besides actin, Myo21 is also present in the flagellar pocket region [15] and in the Myo21^+/−^ cells the impaired intra-vesicular trafficking is accompanied by abnormally large sized flagellar pockets [17]. Further, the data presented here clearly show that in Myo21^+/−^ cells the division of the flagellar pocket slows down, which coincided with the absence of actin-Myo21 co-distribution in this region. These observations suggest that together with actin, Myo21 motor protein is also required to generate force enough to separate the flagellar pockets of dividing daughter cells.

In *Leishmania*, cytokinesis begins as the furrow ingresses between the two (new and old) flagella at the anterior end. As division proceeds, the furrow progresses towards the posterior end, and the two daughter cells, which are placed with their flagella in opposite directions, are eventually separated by abscission [25]. In *Trypanosoma*, the mechanisms and proteins involved in cytokinesis are relatively well studied [9,26]. However, in *Leishmania,* only a few proteins, which are involved in cytokinesis, have been identified, but mechanistic details are still obscure. In *T. brucei*, Aurora B-kinase (AUK1) and Polo-like kinase (PLK) have been implicated in cytokinesis and AUK1 has been shown to regulate cytokinesis initiation, furrow ingression, and abscission [27–30]. In *Leishmania*, coronin distributes at the posterior end during cytokinesis and interacts with dynamic microtubules through kinesis K39, a microtubule-based motor protein [31]. More recently, a homologue of Aurora B kinase in *L. donovani* has been implicated in mitosis and cytokinesis [32].

In trypanosomatids, cell morphogenesis and cytokinesis are closely linked with flagellar biogenesis [33,34]. Ablation of flagellum formation in *Trypanosoma* produces short and immotile cells that are unable to complete cytokinesis, indicating a role of the flagellum in defining the site of cleavage furrow initiation as well as to direct its progression during cytokinesis [35]. Also, flagellar beating provides physical forces required for cytokinesis in *Trypanosoma* [33,34]. Since flagellum in *Leishmania* emerges freely from the cell body, it might not serve the role similar to *Trypanosoma* flagella [35]. This is evident in the case of *L. Mexicana* mutants for an isoform of the dynein (*DHC2.2*), where a shorter flagellum phenotype is not accompanied by any growth defect [36]. Thus, it is less likely that impaired flagellum assembly caused the observed arrest of cell division in Myo21 depleted cells. This argument is supported by the report on *T. brucei* mutants, where depletion of class I myosin does not affect flagellar biogenesis, but still, these cells are defective in cytokinesis [36,37].

The corset microtubules are tightly associated with the subpellicular membrane in the trypanosomatids [38]. Further, actin has been shown to associate with the subpellicular microtubules [39]. However, the physiological significance of actin within the membrane cytoskeleton is still ambiguous. From the results presented here, it may be inferred that Myo21 through actin associates with the subpellicular cytoskeleton, which might be crucial for morphogenesis and cytokinesis. Indeed, the altered morphology of Myo21^+/−^ cells suggests that depleted Myo21 levels are coupled to an impaired cytoskeleton remodeling.

Finally, in trypanosomatids, mitotic checkpoints are not triggered by the defects in cytokinesis, due to which repeated DNA replication results in multinucleated cells under cytokinesis arrest [25]. However, we did not encounter any multinucleated cells in the Myo21^+/−^ cell population despite a cytokinesis block, indicating that karyokinesis might have also been arrested in these cells.

## Acknowledgements

CMG acknowledges the present (HS Subramanya) and the past director (N Yathindra) for providing the facilities and financial support to continue research activities at IBAB. RB would like to acknowledge Dr. S. Thiyagarajan for helpful suggestion as and when required and Mrs. Bindu Ambaru for her help in qPCR.

## Supporting Information

**Table S1.**
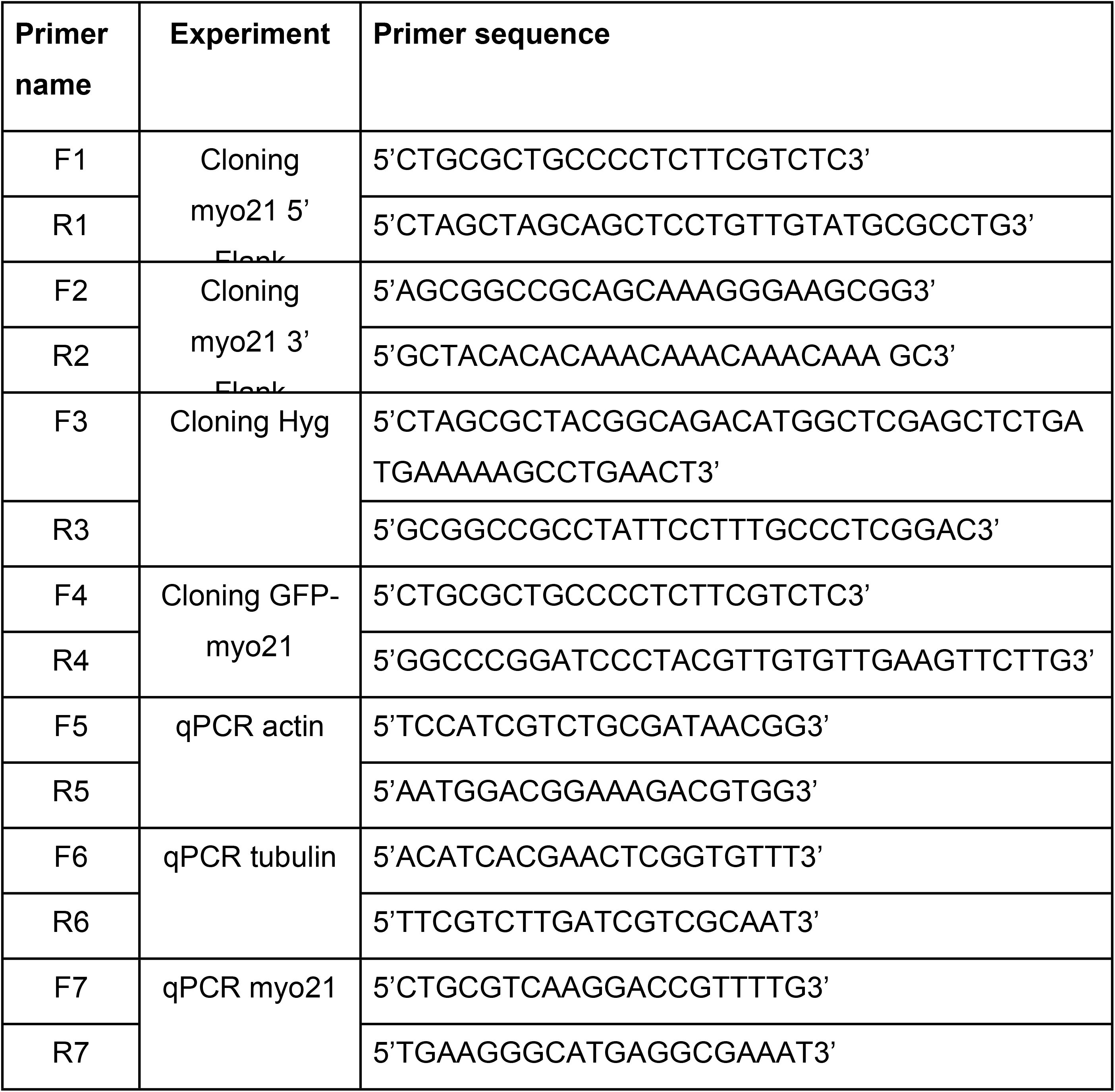
List of primers used in the study.

**Fig. S1:** The original, uncropped and unadjusted images underlying all blots and gels.

**Fig. S2: Cloning of Myo21 deletion cassette.** (A) Schematic representation of strategy for *MYO21* replacement in the *Leishmania* genome. Hygromycin (Hyg, hygromycin phosphotransferase gene) cassette with 5’ FLK (flanking region) and 3’FLK of *Myo21* gene was generated by cloning. (B) Agarose gel images showing B1) PCR amplified Myo21 5’FLK indicated by an arrowhead, B2) PCR amplified Myo21 3’FLK indicated by an arrowhead and B3) Clones (1 & 2) of deletion construct subjected to restriction enzyme digestion to confirm the release of the deletion cassette. Lr: DNA ladder. The arrowhead and asterisk indicate released Hyg cassette (~3kb) and digested vector (~2.8 kb), respectively.

**Fig. S3: Validation of Myo21^+/−^ mutants by qPCR and western blotting.** (A) Western blot analysis showing depletion of Myo21 protein in four independent clones of Myo21^+/−^, as compared to Myo21^+/+^ *Leishmania* cells. Equal cell lysates (equivalent to 10^7^ cells) were loaded, Myo21^+/+^: wild type, Myo21^+/−^: Myo21 heterozygous mutants. Mr: protein molecular weight markers. A1-blot probed with anti-Myo21 antibodies and A2-blot probed with anti-actin antibodies. The asterisk indicates the Myo21 band (~115 kDa). One of the clones was subjected to quantitative expression analysis (B) qPCR analysis showing reduced Myo21 transcript level in Myo21^+/−^ mutants, as compared to Myo21^+/+^ cells. Values shown are the mean of three experiments performed independently ± S.D. ***P-value being <0.0001. (C) Western blot analysis of the Myo21^+/−^ clone. Equal cell lysates (equivalent to 10^7^ cells) were loaded for Myo21^+/+^, Myo21^+/−^ & Myo21^+/−comp^. Mr: protein molecular weight markers. C1-blot probed with anti-Myo21 antibodies and C2 - blot probed with anti-actin antibodies. The asterisk indicates the Myo21 band (~115 kDa), arrowhead indicates Myo21-GFP (~142 kDa). (D) Western blots of three independent experiments were quantified using GelQuant.net software, normalized to actin and the percentage fold change in Myo21 levels has been presented in the bar diagram. Quantification revealed ~70% reduction in Myo21 expression levels in Myo21^+/−^ cells, as compared to Myo21^+/+^ cells. Values shown are the mean of three independent experiments ± S.D. ***P-value <0.0001. Statistical analysis was done by the ANOVA test and a p-value was considered significant if ≤0.05.

**Fig. S4: Cell cycle progression analysis of Myo21^+/+^, Myo21^+/−^ and Myo21^+/−comp^ cells by flow cytometry.** Each cell type was synchronized at the G1/S phase border by hydroxyurea (HU) treatment and after the release of the block; cells were aliquoted at an interval of two hours, processed and stained with PI (propidium iodide). Data were acquired on a flow cytometer and analyzed for distribution of cells in G1, S and G2/M phases, using ModFit software, presented here in histograms. Each data set is representative of three independent experiments and for each sample, 20,000 events were collected. Arrows indicate the position of G1, S and G2/M phases in the histograms. For each cell type, the S phase peaked at the 4^th^ hour, followed by the G2//M phase maxima at the 6^th^ hour. Myo21^+/+^ and Myo21^+/−comp^ cells reappear in the G1 phase by the 8^th^ hour. However, for Myo21^+/−^, the G1 phase maxima reappeared by the 12^th^ hour. Whereas, the G2/M phase has higher percentages of cells for a prolonged time, revealing a delayed progression through the G2/M phase in Myo21^+/−^ mutants.

